# Antibiofilm peptide incorporated PHBV-PLLA nanofibrous mat for wound infection treatment and healing

**DOI:** 10.64898/2026.06.13.732075

**Authors:** Nanditha Chundayil Kalathil, V Reshma Aravind, G.S. Vinod Kumar

## Abstract

Tissue regeneration using bioactive biomaterials has made great progress in the field of wound healing. Biopolymers play a cardinal role in regenerative medicine by providing safe, biocompatible and bioresorbable support. The electrospinning fabrication technique has been used in creating suitable wound care materials. PHBV and PLLA are FDA approved polymers having important applications in biomedical field. In this study, to increase the wound healing potential, PHBV was functionalized with –COOH group and electrospun nano-fibrous mat was produced using PHBV-COOH and PLLA blended solution. Antibiofilm peptide (IDR-1018) with immunomodulatory activity was incorporated into the blended solution to improve infected wound treatment by actively fighting against bacterial infections. Furthermore, in-vitro experiments including cell cytotoxicity assay and scratch wound healing assay were done to evaluate the potential of the synthesized bioactive nanofibrous mat as a potential wound management aid.

## 1. Introduction

Chronic wounds are a growing socio-economic issue. Over one third of the traditional wound healing practices fail to heal chronic wounds and effectively address wound infections. Regenerative medicine technologies have great potential to surpass these problems. One of such successful approach of this technology is the creation and use of three-dimensional scaffold as extracellular matrix analogs that mimic the natural extracellular matrices. These include customizable nano-fiber matrices incorporating novel materials such as peptides, proteins, small molecules etc. [1]. The polymer-based electrospun nano-fibrous scaffolds play prominent role in wound healing. They are excellent skin scaffolds because of their ability to protect the wound area from infection, to prevent fluid and protein loss from wound bed by proper exudate removal and to promote cell adhesion and proliferation. It is due to their increased surface area for cellular interaction with individual fibrils, they exhibits improved cell adhesion, proliferation and migration [2]. To obtain nano-fibrous scaffolds that would match the required properties of wound healing materials, selection of appropriate polymer is very important. The electrospun nano-fibers utilize a range of biopolymers with natural origin such as collagen, chitosan, gelatin, hyaluronic acid and that of synthetic origin, such as poly(hydroxyalkanoate), poly(caprolactone), poly(lactic acid), poly(glycolic acid), and polyurethane [3]. Among them, Poly(hydroxyalkanoates) (PHA) are polyesters produced by micro-organisms under unbalanced growth conditions. PHA are generally biodegradable, with good biocompatibility, making them attractive tissue engineering biomaterials. Some of the widely used PHA include poly 3-hydroxybutyrate (PHB), copolymers of 3-hydroxy butyrate and 3-hydroxy valerate (PHBV), poly 4-hydroxy butyrate (P4HB), copolymers of 3-hydroxy butyrate and 3-hydroxy hexanoate (PHBHHx) and poly 3-hydroxy hexanoate[4].

Poly(3-hydroxybutyrate-co-3-hydroxyvalerate) (PHBV), a biodegradable and biocompatible polyester formed by bacterial synthesis, is a promising scaffolding material for tissue engineering applications. The combination of inherent properties of PHBV and the benefits of fibrous topography could offer a better performance as skin scaffolds. The oxygen permeability, nano-fibrous topography, mechanical property and superior cell-scaffold interaction contributes to the higher proliferation on PHBV scaffold. The porosity of the scaffold coupled with the inherent oxygen permeability property of PHBV would facilitate nutrient diffusion and gas exchange that is imperative for cell survival in culture. Human keratinocytes (HaCaT) adhesion, proliferation and gene expression were higher on PHBV nanofibers [2]. PLA is a linear aliphatic thermoplastic produced by the fermentation of corn dextrose. It is biodegradable and characterized by a high tensile strength and modulus but low ductility. Poly(L-lactic acid (PLLA) is synthesized from the polymerization of the L-form of lactic acid. The utilization of PHBV alongside PLLA improves the biocompatibility and wettability of electrospun polymer scaffolds used for tissue engineering over neat electrospun PLLA. The biocompatibility of PLLA and PHBV, combined with the inherent porosity of electrospun scaffolds, make the polymer blend well suited to electrospinning [5].

Wound infections and biofilm formation is a major issue in chronic wound treatment as it impedes the natural healing process. Promotion of wound healing in such cases can be achieved by the use of antimicrobial and antibiofilm peptides. Innate Defence Regulator peptides (IDR peptides) are synthetic immunomodulatory versions of natural host defence peptides (HDP). In the absence of direct antimicrobial activity, IDRs mediate protection against bacterial challenge. It thus represents a novel approach to anti-infective and anti-inflammatory therapy. IDR peptides possess the ability to enhance wound healing. Studies have reported that IDR-1018, a synthetic peptide, helps in faster wound closure in a concentration-dependent manner. IDR-1018 also enhances re-epithelialization by promoting keratinocyte proliferation. It works by modulating immune responses [6].

In this study, we have developed an electrospun fibrous mat as a medium for delivering antibiofilm peptide for enhanced wound healing activity. To increase wound healing properties and promote peptide adsorption, PHBV was functionalized with –COOH group. PHBV and PLLA were blended together in order to increase stability and electrospun to form a nano fibrous mat. IDR-1018 peptide was incorporated into the nanofibers as a bioactive material since it has been reported that the peptide acts as a potent anti-microbial and antibiofilm agent and improves wound healing efficiency. The functionalized nanofibrous mat was characterized and in-vitro cytotoxicity and scratch wound assay were conducted to determine the efficacy of the developed wound healing aid.

## 2. Materials and Methods

### 2.1 Materials

Poly (3-hydroxybutyrate-co-hydroxyvalerate) (PHBV), hydroxyl valerate content = 12 mol%), 3-(4,5dimethylthiazol-2-yl)-2,5-diphenyltetrazolium bromide (MTT), 4-(dimethyl amino) Pyridine (DMAP), Phosphate buffered saline (PBS), Triethyl Amine (TEA), Hydroxymethyl-phenoxyaceticacid (HMPA), N-hydroxybenzotriazole (HOBT), N-Methyl Imidazole (Sigma-aldrich), Ninhydrin, Trifluroacetic acid (TFA), Triisopropylsilane (TIS) and Succinic anhydride were procured from Sigma-Aldrich. Tentagel-S-NH2 resin (0.29 meq/g), Fmoc protected amino acids from Peptide International,Inc. N,N,N’,N’,Tetramethyl-O-(1H-benzotriazol-1-yl) uranium hexafluorophosphate (HBTU) from Spectrachem. N,N-diisopropylethylamine (DIEA), 1-(mesitylene-2-sulphonyl)-3-nitro-1H-1,2,4-triazole (MSNT) from Nova Biochem. Analytical grade chloroform, Dichloromethane (DCM) and N,N-dimethylformamide (DMF) from Merck, Piperidine, Diethyl ether were of analytical grade, poly (L-lactic acid) (PLLA, MW=50,000) purchased from Polysciences. Dulbecco’s modified eagle medium (DMEM), Fetal bovine serum (FBS) was obtained from Invitrogen.

### 2.2 Peptide synthesis

Innate Defence Regulator peptide (IDR-1018) (VRLIVAVRIWRR-NH2) [6] peptide was synthesized by solid phase peptide synthesis using the standard Fmoc-Strategy [7]. TentaGel™–S–NH2 resin was linked to HMPA linker by HBTU/HOBT activation. The peptide synthesis was then carried out using the standard Fmoc procedures. After synthesis, the peptide was manually cleaved using Trifluoroacetic acid (TFA), Triisopropylsilane, and water (95:2.5:2.5). The peptide was then precipitated with diethyl ether, followed by 15–25 washes with diethyl ether. The peptide was dissolved in minimum amount of water, lyophilized and stored at −20°C. The peptide identity was verified by matrix-assisted laser desorption ionization time-of-flight mass spectroscopy.

### 2.3 Functionalization of PHBV

Carboxyl group (COOH) modification of PHBV was done under inert conditions. 20g PHBV was dissolved in 100ml chloroform by gentle stirring in a 250ml RB using a hotplate with magnetic stirrer for 90 minutes in an oil bath of 1000C. The mixture was then added with 220mg 4-(dimethyl amino) pyridine (DMAP), 240μl Triethyl amine and 5g Succinic anhydride. RB containing the mixture was kept in magnetic stirrer for 48 hours at room temperature. The viscous solution was dried in rotary evaporator to remove solvent. Solution was filtered through syringe filter [(MILLEX-HA)-MCE 0.45μm filter] and was precipitated in ice cold ether. Filtrate was kept in ice and colour of the precipitate was noted. Top layer is removed and ether is added. The process is repeated for 2-3 times for purification. It is then kept for drying and stored in desiccator. The COOH-functionalization of PHBV was characterised using Fourier-transform infrared spectroscopy (FTIR).

### 2.4 Preparation of electrospun PHBV-PLLA nanofibers

A preliminary investigation was first conducted to determine the optimal ratio of the concentration for the electrospinning solution, and the solution concentration was fixed at 30 wt%. The optimal PHBV: PLLA blend ratio was fixed at 5: 1. PHBV and PLLA solutions were prepared by dissolving in chloroform separately. 0.0030g IDR-1018 peptide was dissolved in 1ml methanol. The three solutions were then mixed together using magnetic stirrer, for making a blend solution. Electrospinning was performed in HO-NFES-043 Halmarc Nano Fiber Electrospinning Unit. For the principle electrospinning trials, a total polymer content of 30 w/v, needle gauge of 0.55 mm and collector distance of 10 cm were held constant while the blend ratio was varied. For each blend ratio, the applied voltage and solution feed rate were optimized to achieve continuous spinning and smooth fibers, with the latter checked by optical microscope. A total of 20 ml of solution was electrospun onto the collector (collection time varied based on cleaning frequency of the needle and feed rate). To evaporate any residual solvent, electrospun fibers were kept under a fume hood for 24 h prior to characterization and storage.

### 2.5 Characterization of electrospun fiber mats

Optical microscopy was used for the immediate study of the fiber integrity and mat formation for optimization of the electrospinning voltage and feed rate for each polymer blend ratio. A glass microscope slide was placed on the aluminium collection plate and allowed to collect fibers for 1−2 min. The microscope slide was then viewed under the optical microscope at 20X magnification to check for the formation of fibers and the presence of beads. Feed rate and voltage were adjusted for each solution until continuous fibers with the least amount of beading were formed.

The nanofiber mat was analyzed by Differential Scanning Calorimetry (DSC). DSC thermograms were obtained using Pyris TM DSC 6000 PerkinElmer, Waltham, MA. The sample was loaded in standard aluminium pans and was heated from 0 °C up to 200 °C at a rate of 10 °C/minute under constant nitrogen purging of 20 ml/minute. An empty pan was used as a reference. The developed nanofiber was also characterized using FTIR. The spectra were recorded between 600 and 4000 cm^−1^ wave number range using Nicolet TM 5700 spectrometer.

### 2.6 In-vitro Tests

Human keratinocyte HaCaT cell line was purchased from National Centre for Cell Science, Pune. The cells were grown in DMEM supplemented with 10% FBS and 1% antibiotics.

#### 2.6.1 Cytocompatibility study

The cell viability of the nanofiber scaffold was assessed by MTT (3-(4, 5-dimethylthiazol-2-yl)-2, 5-diphenyltetrazolium bromide) reduction assay. The nanofiber mat was cut into a pre-determined size and placed in a 24 well plate. The samples were sterilized before cell seeding and then washed with PBS three times and incubated with DMEM medium for 1 h. The cells seeded on the scaffolds at a density of 5 × 104 cells per ml and incubated at 37 °C. After 24, 48 and 72 h, MTT was added to the wells and further incubated for 4 h. Absorbance was measured at 570 nm using Biorad iMark™ Microplate Absorbance Reader. The cell viability was evaluated from triplicate sample results. Percentage viability was calculated using the formula: [(Avg. OD of test / Avg. OD of control) × 100].

#### 2.6.2 Scratch Wound Assay

HaCaT cells were grown in 12 well culture plates and were then scratched with 2.5μL pipette tip and a uniform cell-free zone was created in each well. The wells were washed with PBS to remove cellular debris. Media treated with nanofiber mat were added to the wells and incubated for 24 hours. The cell culture wells were observed at 0 hour and 24 hour time points after injury. Images were taken and the wound closure diameter (µm) between wound edges was calculated with computer-assisted analysis system (Olympus CellSens). The wound closure percentage was calculated using the formula: [% Wound closure = (Pre-migration) diameter − (Migration) diameter / (Pre-migration) diameter ×100]

## 3. Results and Discussion

Microbial infection and biofilm formation is a major problem associated with chronic wounds. One of the bases for the treatment of wound infections is the use of antibiotics. But the effectiveness of antibiotics has become restricted due to an increase in bacterial antibiotic resistance. One promising set of compounds which can be used for infected wound treatment is cationic antimicrobial peptides which represent a good template for a new generation of antimicrobials. They kill both Gram negative and Gram positive microorganisms rapidly and directly, work against common clinically-resistant bacteria such as methicillin-resistant Staphylococcus aureus (MRSA) and vancomycin resistant Enterococcus (VRE), show a synergistic effect with conventional antibiotics, and can often activate host innate immunity without displaying immunogenicity [8]. Promotion of complete healing in infected wounds can thus be achieved by the delivery of known antimicrobial and antibiofilm peptide to the wound site. IDR-1018 (HOOC-VRLIVAVRIWRR-NH2) is a reported antibiofilm peptide which also possesses wound healing ability. In this study, the peptide IDR-1018 with twelve amino acid sequence, containing poly-valine in its C terminal was synthesized by following a standard protocol of solid phase peptide synthesis technique. MALDI characterization of the crude solution confirmed the presence of IDR 1018 peptide. The cell viability potential of the peptide alone in HaCaT cells were analyzed for concentrations ranging from 5μg/ml to 100μg/ml [Fig.1] and was found that the peptide showed good viability percentage. Based on the above result, IDR1018 peptide was selected as the bioactive agent for incorporation to nanofiber mat as infected wound dressing.

**Fig. 1.**
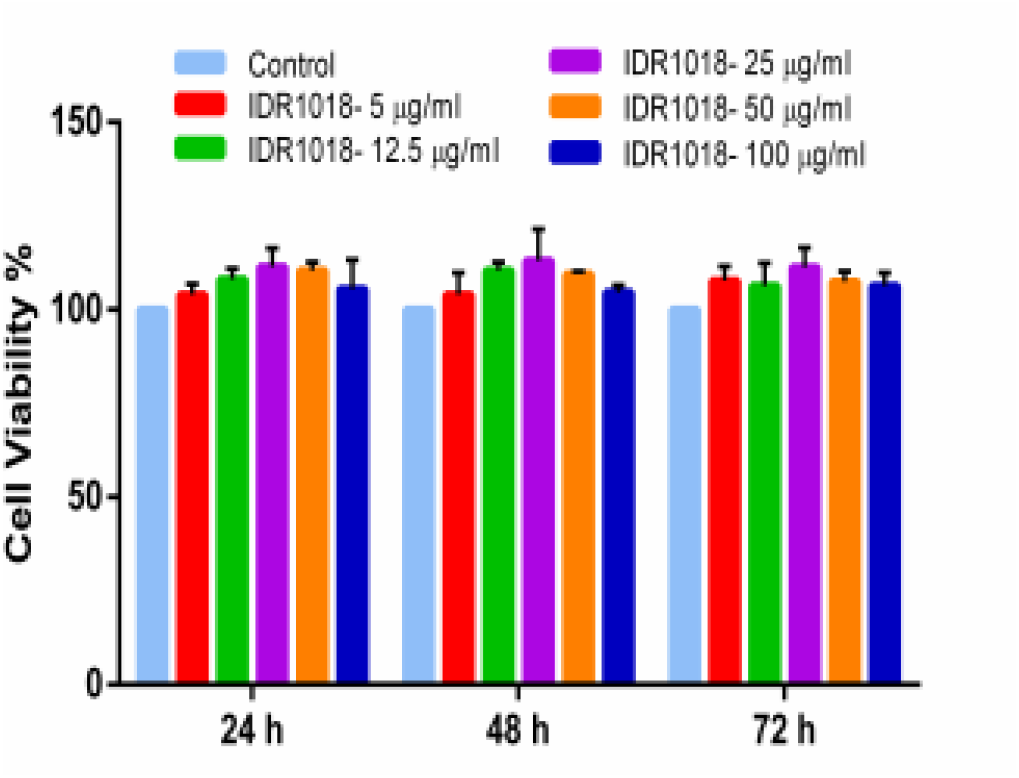
MTT (3-(4, 5-dimethylthiazol-2-yl)-2, 5-diphenyltetrazolium bromide) Assay of IDR1018 peptide at different concentrations showing their cytocompatibility in HaCaT cell line at 24, 48 and 72 hours.

Biopolymers are more effective wound healing agents than traditional wound dressings due to their potential drug delivery property through controlled release and also high biocompatibility along with application easiness [9]. Electrospinning technique allows the production of ultra-fine fibers that would mimic the structure and biological functions of the natural extra cellular matric (ECM) which is required for the restoration of tissue function. Carboxylic group functionalized PHBV and PLLA solutions mixed with IDR 1018 peptide will help in the slow release of the antibiofilm agent.

The developed electrospun nanofiber mat was characterised using Fourier transform infrared spectroscopy (FTIR) analysis and Differential Scanning Calorimetry (DSC) analysis to understand and confirm the presence of modifications and thermal stability. As represented in the infrared spectrum of PHBV-COOH [Fig.2(a)], the absorbance peaks at 2978, 2933, and 1720 cm^−1^ were asymmetric and symmetric stretching vibration of CH3 and stretching vibration of C=O of PHBV, respectively. A very strong and sharp absorption band at 1720.69 cm−1 observed in the FTIR spectrum of COOH modified PHBV is attributed to the C = O stretching mode. The FT-IR spectra of the PHBV COOH/PLLA blend [Fig.2(b)], with characteristic absorbance bands were recorded in the regions 1800 – 1600 cm^−1^ and 1000 – 700 cm^−1^. The carbonyl absorption bands in the range of 1760 to 1730 cm^−1^ in PHBV COOH/PLLA blend is also characteristic in the spectrum, which is usually based on the blending ratios of PHBV and PLLA, mostly as a result of change in local molecular environment during crystallization

**Fig. 2.**
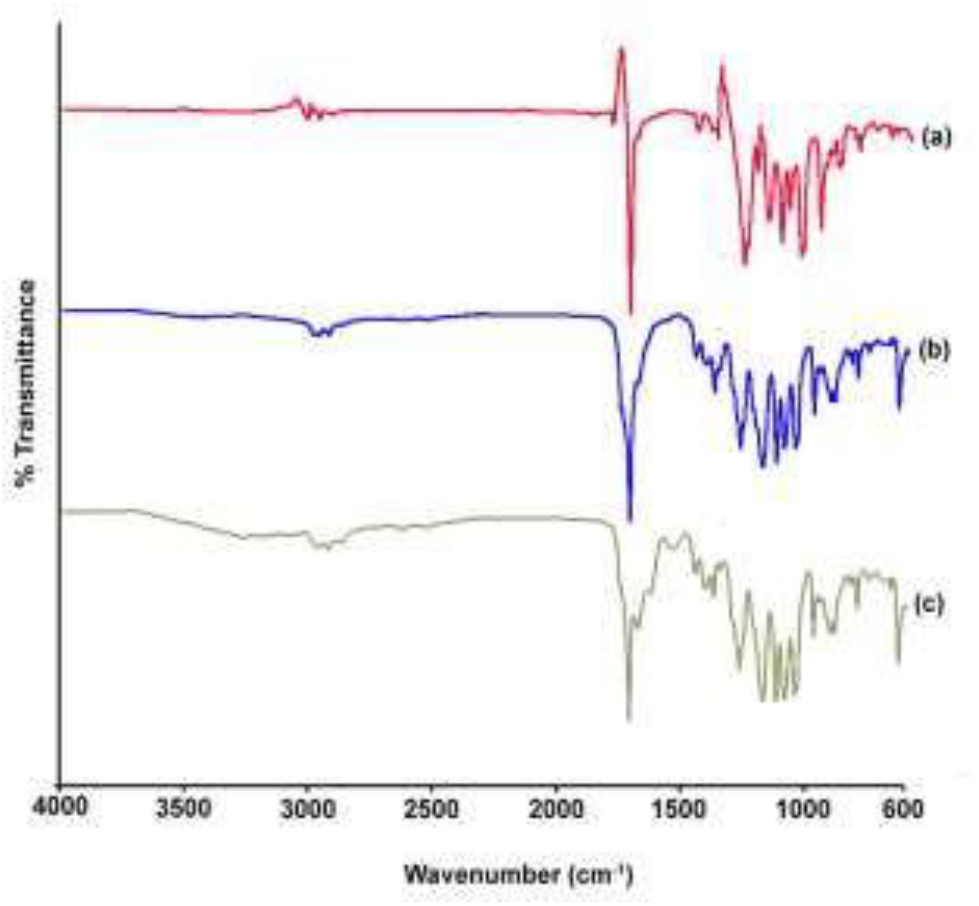
Fourier transform infrared spectroscopy (FTIR) analysis spectra of (a) PHBV-COOH, (b) PHBV-COOH/PLLA and (c) IDR1018/PHBV-COOH/PLLA

Accordingly, the addition of PLLA to PHBV does not change the overall helical structure of the blends, but alters the helical structure from place to place, leading to a decrease in crystallinity resulting from the formation of a number of amorphous parts. The presence of the peptide IDR1018 in the electrospun blend of PHBV COOH/PLLA was confirmed using FTIR [Fig.2.(c)]. Apart from the characteristic peaks of PHBV COOH/PLLA blends, the absorbance peak at 1685 cm-1and 1545 cm-1 represent the C=O stretch and N-H bending vibrations in amide bonds. Presence of a broad absorbance band in the range of 3000 to 3500 cm-1 represents the N-H stretching vibration characteristic to amide bonds, thus confirming the presence of the peptide.

In the Differential Scanning Calorimetry (DSC) spectrum, the miscibility of the pure polymers in a blend can be determined by observation of the glass transition temperature of the blend. A single composition-dependent glass transition temperature for a polymer blend, located between the glass transition temperatures of the polymers, is an indication of polymer miscibility.

A bimodal endothermic melting peak for PHBV COOH was observed, mostly as the result of the modification process [Fig.3 (a)]. The cause of multiple peaks may be due to the formation of heterogeneous crystals and their melting, recrystallization, and remelting [10]. The melting peaks of the electrospun PLLA and PHBV COOH blend were observed at 170 and 135°C [Fig.3 (b)]. The obtained results indicate some degree of engagement between PHBV COOH and PLLA chains in the fibers or partial miscibility that may be caused by high shear and elongation during the spinning process.

**Fig. 3.**
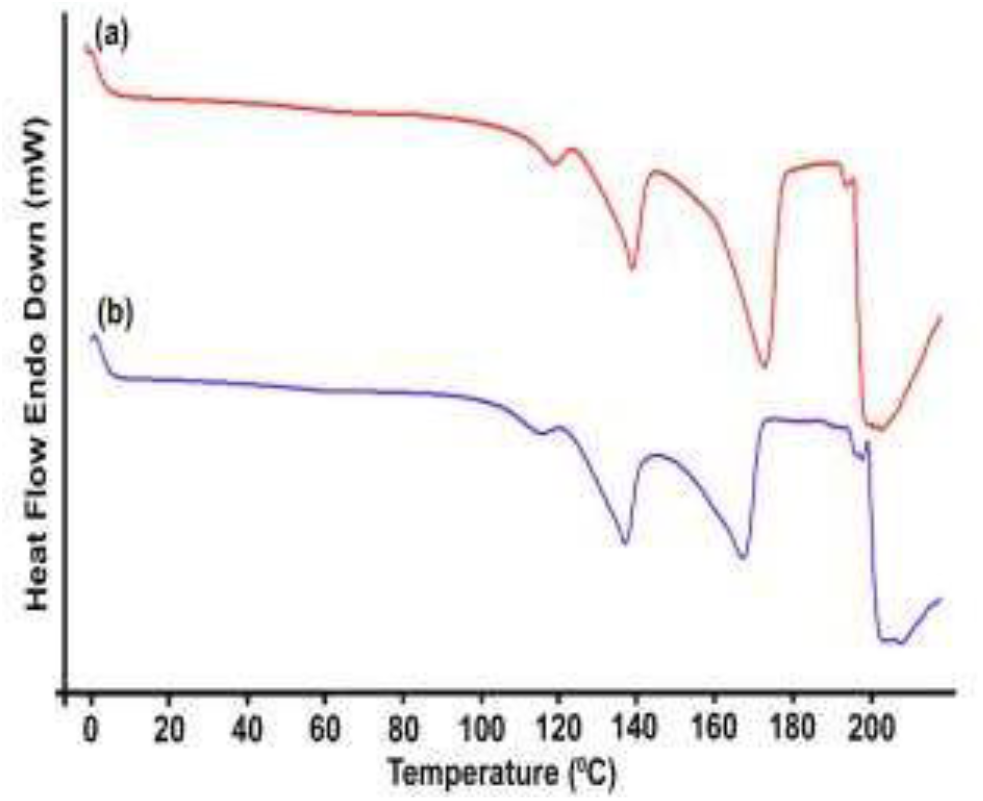
Differential Scanning Calorimetry (DSC) spectrum of (a) PHBV-COOH and (b) PHBV-COOH/PLLA

The biocompatibility of antibiofilm peptide carrying PHBV-COOH/PLLA as a wound management aid was evaluated against in vitro cultures of HaCaT cells [Fig.4]. The peptide carrying nanofiber mat showed no cytotoxicity on HaCaT cells and the slow release of peptide maintained their bioactivity even when incorporated into the nanofiber as shown by the increased cell viability when compared to nanofiber alone group. It was found that the nanofiber mat without the bioactive peptide when compared to the control did not exhibit much cytotoxicity.

**Fig. 4.**
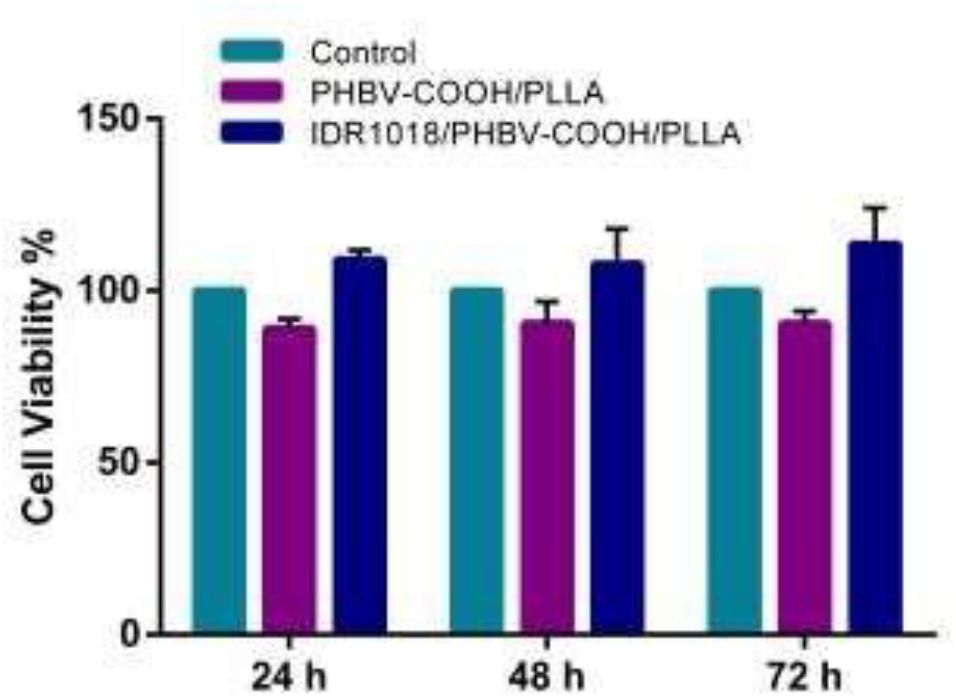
MTT (3-(4, 5-dimethylthiazol-2-yl)-2, 5-diphenyltetrazolium bromide) Assay of PHBV-COOH/PLLA and IDR1018/PHBV-COOH/PLLA in HaCaT cell line at 24, 48 and 72 hours.

Re-epithelialization is an important process of wound healing and is identified by increased keratinocyte proliferation and migration over the wound area [11]. The in-vitro scratched wound healing data [Fig.5(a)] showed that the peptide carrying nanofiber treated sample showed significant healing effects within 24h of treatment. The percentage wound closure [fig. 5(b)] by the nanofiber and IDR1018/ PHBV-COOH/PLLA nanofiber mat treated cells displayed much better wound healing action when compared to the control. The wound closure data obtained from the scratched wound healing assay gives an idea about the efficiency of the IDR1018 peptide entrapped PHBV-COOH/PLLA nanofiber as a potential wound management aid.

**Fig. 5(a).**
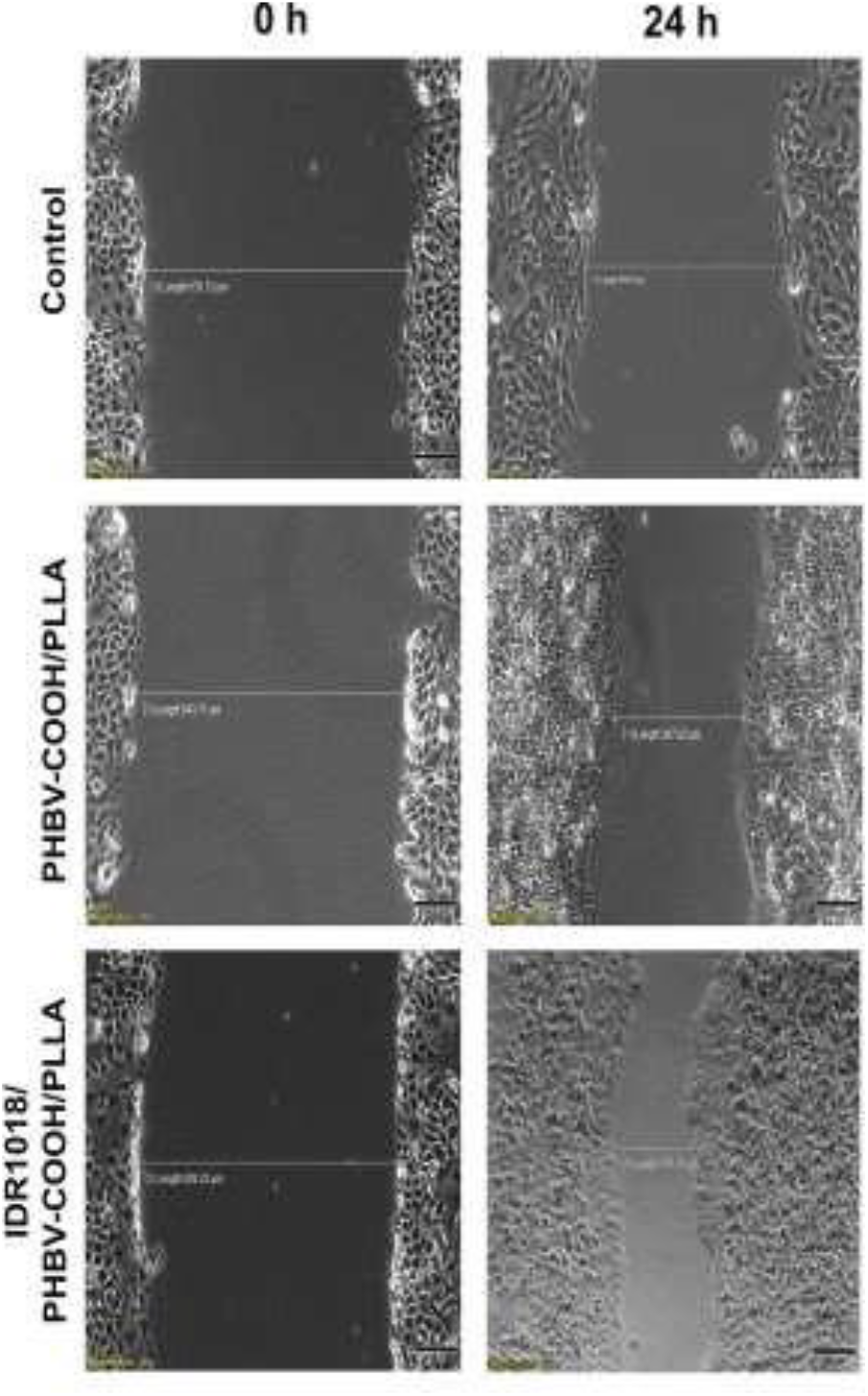
Scratched wound healing assay result in HaCaT cell line

**Fig. 5(b).**
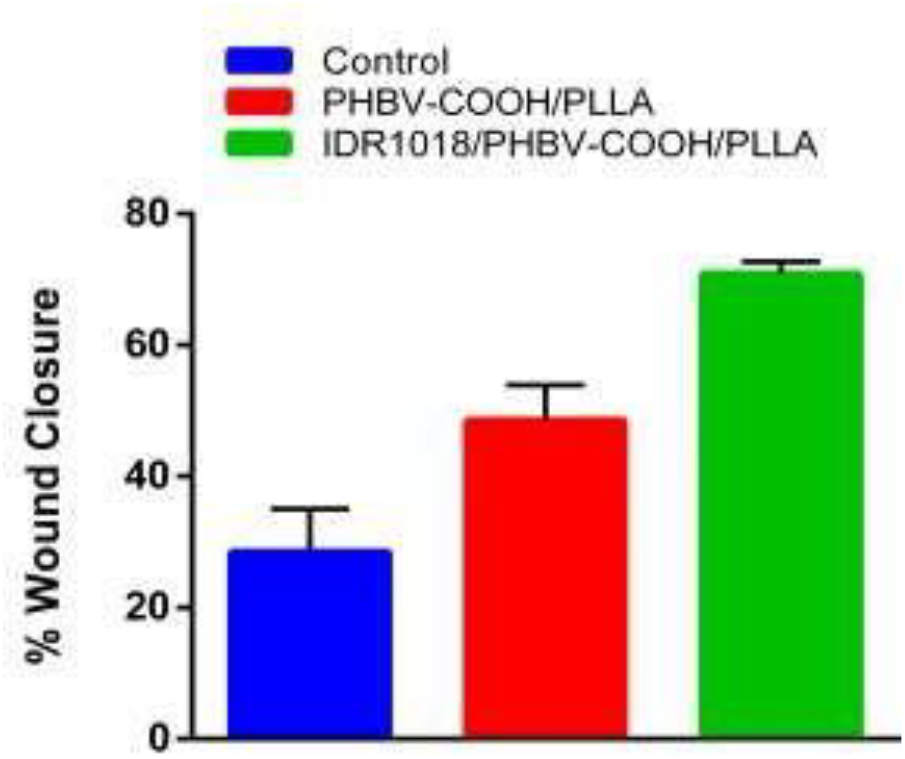
Percentage wound closure in HaCaT cells treated with PHBV-COOH/PLLA and IDR1018/PHBV-COOH/PLLA

## 4. Conclusion

To overcome the drawbacks of the traditional wound treatments involving infected wounds, an efficient electrospun mat incorporated with antibiofilm peptide was developed. FDA approved polymers PHBV and PLLA were blended together along with bioactive peptide IDR-1018 for enhanced wound healing activity. PHBV was functionalized with -COOH for increasing wound healing property and peptide adsorption. To overcome the instability of PHBV fibrous mat, it was electrospun by blending with PLLA and characterized by FTIR and DSC. In vitro studies were done by analyzing the cell cytotoxicity and by scratched wound assay and the antibiofilm peptide incorporated nano-fibrous mat was found to be efficient as a potential wound management aid.

## Acknowledgment

The authors are grateful to the Department of Biotechnology, New Delhi, for providing the financial assistance, the University Grants Commission, New Delhi, for the Junior Research Fellowship to Nanditha C.K, and University of Kerala for PhD registration.

